# Breast tumor-associated metalloproteases restrict reovirus oncolysis by cleaving the σ1 cell-attachment protein, and can be overcome by mutation of σ1

**DOI:** 10.1101/742478

**Authors:** Jason Fernandes, Francisca Cristi, Heather Eaton, Patricia Chen, Sarah Haeflinger, Isabelle Bernard, Mary Hitt, Maya Shmulevitz

## Abstract

Reovirus is undergoing clinical testing as an oncolytic therapy for breast cancer. Given that reovirus naturally evolved to thrive in *enteric* environments, we sought to better understand how *breast tumor* microenvironments impinge on reovirus infection. Reovirus was treated with extracellular extracts generated from polyoma virus middle T-antigen-derived mouse breast tumors. Unexpectedly, these breast tumor extracellular extracts inactivated reovirus, reducing infectivity of reovirus particles by 100-fold. Mechanistically, inactivation was attributed to proteolytic cleavage of the viral cell attachment protein σ1, which diminished virus binding to sialic acid-low tumor cells. Among various specific protease class inhibitors and metal ions, EDTA and ZnCl_2_ effectively modulated σ1 cleavage, indicating that breast tumor-associated zinc-dependent metalloproteases are responsible for reovirus inactivation. Moreover, media from MCF7, MB468, MD-MB-231 and HS578T breast cancer cell lines recapitulated σ1 cleavage and reovirus inactivation, suggesting that inactivation of reovirus is shared among mouse and human breast cancers, and that breast cancer cells in by themselves can be a source of reovirus-inactivating proteases. Binding assays and quantification of sialic acid (SA) levels on a panel of cancer cells showed that truncated σ1 reduced virus binding to cells with low surface SA. To overcome this restriction, we generated a reovirus mutant with a mutation (T249I) in σ1 that prevents σ1 cleavage and inactivation by breast tumor-associated proteases. The mutant reovirus showed similar replication kinetics in tumorigenic cells, equivalent toxicity as wild-type reovirus in a severely compromised mouse model, and increased tumor titers. Overall, the data shows that tumor microenvironments have the potential to reduce infectivity of reovirus.

**SIGNIFICANCE:** We demonstrate that metalloproteases in breast tumor microenvironments can inactivate reovirus. Our findings expose that tumor microenvironment proteases could have negative impact on proteinaceous cancer therapies such as reovirus, and that modification of such therapies to circumvent inactivation by tumor metalloproteases merits consideration.

## Introduction

Mammalian orthoreovirus serotype 3 (reovirus, T3D) has been known for some time to possess oncolytic activity and is being investigated as a treatment against numerous cancers (1–3). In immunocompetent hosts, reovirus is asymptomatic and therefore a safe candidate for oncolytic therapy, showing minimal side effects in numerous human clinical trials (4–8). Reovirus oncotherapy shows promise, for example improving overall survival from 10.4 months to 17.4 months when combined with paclitaxel in an ongoing breast cancer trial (9). Nevertheless, reovirus therapy fails to cure the majority of immunocompetent tumor-bearing animals and human patients (10–13), indicating that we must better understand the restrictions to reovirus treatment and ways to improve the potency of this therapeutic agent.

We posited whether the heterogeneity of tumor environments could contribute to differential activity of reovirus among tumors. The natural route of infection for reovirus is predominantly enteric, and consequently reovirus has evolved to be exquisitely well adapted to the human gastrointestinal tract. For example, reovirus evolved a very stable double-layered proteinaceous capsid that withstands harsh conditions of the environment and enteric tract. The outercapsid is composed of structural proteins µ1 and σ3, and a third protein σ1 that protrudes from viral vertices and facilitates cell attachment. The inner capsid ‘core’ is composed of additional structural proteins, transcription factors, and the dsRNA genome (14–16). Importantly, to mediate efficient disassembly of the outercapsid which is required for reovirus to establish infection, reovirus exploits gut proteases chymotrypsin and trypsin. Specifically, these proteases generate infectious subvirion particles (ISVPs) by degrading the outer-most capsid protein σ3 and cleaving the underlying µ1 protein into multiple fragments that promote membrane penetration and entry of reovirus particles into the cell (17–20). Membrane penetration results in disassociation of the remaining µ1 outercapsid fragments and σ1 (21), delivering transcriptionally-competent core particles. In the absence of trypsin or chymotrypsin, reovirus conversion to ISVPs can still occur within endo-lysosomes, mediated by lysosomal cathepsins B and L (22). The dependency of reovirus on gut and lysosomal proteases provoked an important question for reovirus infectivity in tumors; what sort of proteases are in tumor environments and do they support reovirus uncoating?

The tumor microenvironment presents a distinct set of proteases relative to the natural enteric route of reovirus infection. Although chymotrypsin and trypsin are restricted to the gut, tumors are rich in other proteases such as various cathepsins and metalloproteases which have been shown to play important roles in cancer progression (23–25). For example, tumor invasiveness and metastasis require degradation of the extracellular matrix (ECM), which is facilitated by proteolytic pathways involving multiple families of proteases (26, 27). Cathepsin B is classically considered a lysosomal protease, however, extracellular and pericellular translocation of the enzyme was observed in cancers such as breast and brain cancers, and can correlate with poor prognosis (23, 28–31). Cathepsin B is implicated in the activation of other ECM-degradation associated proteases such as urokinase plasminogen activator, which in turn, proteolytically activates numerous metalloprotease downstream targets that contribute to ECM degradation (32). But whether tumor proteases can target reovirus proteins, and whether they are in sufficient concentration to modulate reovirus infectivity, is unknown.

The current study aimed to determine if breast tumor extracellular proteases have any effect on reovirus, and if so, whether the effect is positive, negative or neutral towards reovirus infectivity of tumor cells. We prepared tumor extracellular extracts (T.E.E.) from murine breast tumors to mimic the tumor microenvironment then treated reovirus with T.E.E. under varying conditions. The three key discoveries of our study are (1) T.E.E. contains zinc dependent metalloprotease activity that cleaves the cell attachment protein σ1 and [sometimes] σ3, yet does not produce the beneficial cleavage of μ1C to δ, leading to a 100-fold reduction of reovirus infectivity in many cell types, (2) T.E.E proteases reduce binding of reovirus towards cancer cells that have low sialic acid levels, because cleavage of σ1 removes the JAM-A binding capacity and forces a dependence on sialic acids for attachment, and (3) inactivation of reovirus by breast tumor proteases can be circumvented by mutation of the σ1 protein.

## Results

### Breast tumor extracellular protease(s) cleave reovirus σ1 protein but do not produce ISVPs

The fate of reovirus when exposed to intestinal digestive enzymes is well characterized. Degradation of outer capsid protein σ3 and cleavage of µ1C to δ generates infectious subviral particles (ISVPs) with the ability to directly penetrate membranes, therefore enhancing infectivity (19, 33, 34). For some strains of reovirus, the cell attachment protein σ1 is also cleaved in half, maintaining the sialic acid-binding σ1-tail fragment (σ1N) which permits attachment to the sialic-acid-rich luminal face of intestinal cells. However, with reovirus currently in phase III clinical trials as an oncolytic therapy, it is important to know whether tumors also release proteases that digest reovirus proteins, and if so, whether tumor-associated proteases have positive, negative, or neutral effects on reovirus infection of tumor cells.

To determine the effects of tumor extracellular proteases on reovirus, we generated protease-enriched tumor extracellular extract (T.E.E.) from polyoma virus middle T-antigen-derived mouse breast tumors by diffusion of extracellular content into phosphate-buffered saline (PBS). We then exposed purified reovirus to T.E.E. and monitored reovirus protein processing by Western blot analysis. As a positive control for protease-mediated reovirus uncoating, we also exposed purified reovirus to intestinal extracellular extract (I.E.E.) prepared by flushing murine intestines with PBS. As anticipated, I.E.E. treatment resulted in hallmarks of ISVP formation, including degradation of outer capsid protein σ3, and cleavage of µ1C to δ (Figure 1A). I.E.E. treatment also led to cleavage of σ1 to the σ1N-tail fragment. Despite processing of the outer capsid, the inner core proteins λ1/2 and σ2 remained intact; this was anticipated since the reovirus core becomes transcription-competent and does not further uncoat, but rather synthesizes and expels positive-sense RNA into the cytoplasm during virus replication (35). Importantly, following treatment with T.E.E., σ1 was cleaved to the 22 KDa σ1N fragment. Cleavage of outer capsid µ1, which is a hallmark of ISVP generation, was however limited. Core viral proteins were unaffected by T.E.E. treatment. The effects of T.E.E. were therefore distinct from I.E.E, with T.E.E. causing cleavage of σ1 but failing to generate ISVPs.

**Figure 1:**
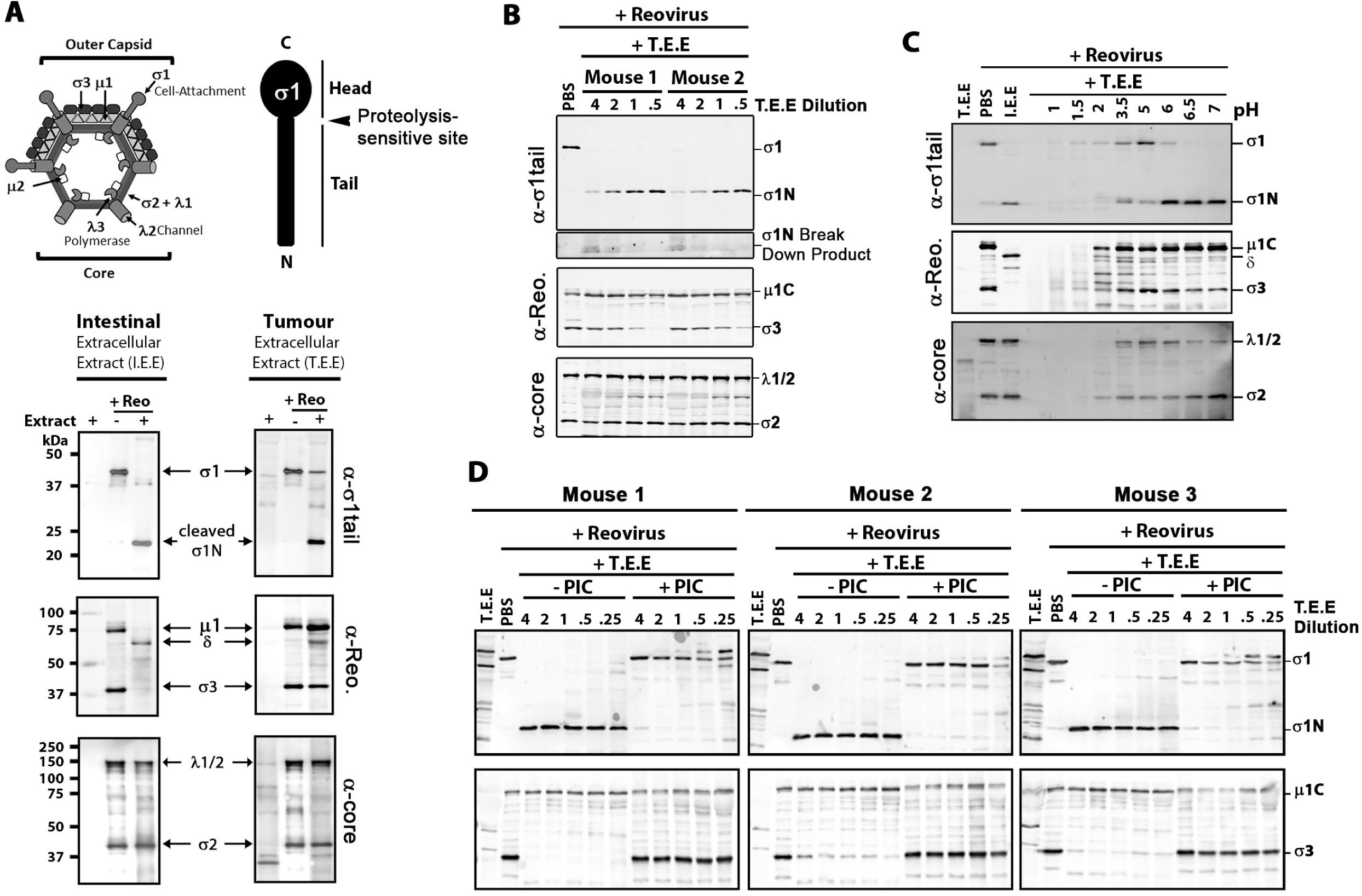
Breast tumor extracellular protease activity cleaves reovirus σ1 protein but does not produce ISVPs. Extracellular contents of murine breast tumors (tumor extracellular extract; T.E.E.) were used to mimic the tumor microenvironment. To mimic the intestinal environment, the protease rich luminal contents were obtained from murine intestines (intestinal extracellular extract; I.E.E.). **A. (TOP)** Diagrammatical depiction of reovirus outercapsid and core proteins, along with a diagram of σ1 tail and head domains. **(BOTTOM)** Reovirus was treated with T.E.E. or I.E.E. at 37°C for 24 hours and viral proteins were subjected to Western blot analysis using antibodies against σ1-tail, outer capsid proteins (polyclonal anti-reovirus serum) or viral core proteins (polyclonal anti-reovirus core serum). **B**. As with A., but the activity of T.E.E. from two additional and independent murine breast tumors was assessed. **C**. Reovirus was treated with T.E.E. under different pH conditions, ranging from pH 1 to pH 7. **D**. Reovirus was treated with T.E.E. from 3 murine breast tumors in the absence or presence of protease inhibitor cocktail (PIC).

Proteolytic processing of reovirus outer capsid proteins can depend on the concentration of proteases, therefore we assessed the fate of reovirus proteins by western blot analysis after exposing purified reovirus particles to increasing amounts of T.E.E. (Figure 1B). Furthermore, to determine if reovirus processing was consistent among distinct T.E.E. samples, we obtained T.E.E. from two independent mouse breast tumor explants. Cleavage of σ1 to σ1N was observed for T.E.E. obtained from both mice, in a concentration dependent manner. Specifically, while all T.E.E. concentrations effectively cleaved σ1 to σ1N, high T.E.E.-to-reovirus ratios caused further degradation of the σ1N 22kDa fragment. While core proteins were unaffected by T.E.E., outer capsid σ3 was degraded under low ratios of T.E.E.-to-reovirus. This could possibly be a consequence of protease-inactivation at high concentrations due to self-cleavage or cleavage by other proteases. Notably, however, under no condition was µ1C processed to δ, suggesting that T.E.E. factors from multiple mice tumors did not yield ISVPs. This is the first time, to our knowledge, that processing of reovirus by T.E.E. has been characterized.

Since breast tumors have slightly acidic environments (36, 37), we hypothesized that incomplete processing of reovirus σ3 or µ1C by T.E.E. could be a result of poorly optimized pH conditions. Therefore, reovirus was treated with T.E.E. at a range of pH conditions (Figure 1C). At highly acidic pH (≤ 2), reovirus particles were degraded completely, as seen by the absence of all viral proteins including core proteins λ1 and σ2. Cleavage of σ1 was most efficient above pH 6, with complete conversion of σ1 to σ1N. Importantly, under no conditions that preserved reovirus core proteins (pH ≥ 2), was µ1C processed to δ. Overall, it seems that the optimal pH for reovirus processing was 6-7, consistent with the environmental pH of most tumors (38, 39).

To confirm that processing of reovirus proteins by T.E.E. was attributed to a protease, reovirus was treated with T.E.E. from three independent mouse tumors but in the presence or absence of a protease inhibitor cocktail (PIC) (Figure 1D). Processing of σ1 and σ3 was strongly impaired in the presence of the PIC at all T.E.E.:reovirus ratios tested, implicating that a secreted protease in mouse breast tumors is involved in cleavage of σ1 and occasional degradation of σ3.

### Breast tumor extracellular protease(s) reduce reovirus attachment and decrease infectivity by 100-fold

Having discovered that T.E.E. cleaves σ1 but does not produce ISVPs, it became important to determine whether this partial processing of reovirus by T.E.E. affects infectivity. Specifically, the cleavage of σ1 could be an important impediment to virus-cell attachment. Normally reovirus σ1 mediates attachment to cells through two domains: the head domain of σ1 mediates high-affinity interaction with junction adhesion molecules (JAM-A), while the tail domain of σ1 supports association with α2,3-, α2,6-, or α2,8-linked sialylated glycans (40). In the intestine, the luminal surface of intestinal cells displays sialic acids (SA) but not JAM-A. Therefore, cleavage of σ1 and removal of the JAM-binding head of reovirus by gut proteases trypsin and chymotrypsin is unlikely to impeded virus binding to the SA-rich and JAM-A-deficient luminal face of intestinal epithelial cells. But JAM-A binding may instead be necessary to mediate attachment at the JAM-rich basolateral side (41–43) of the intestinal epithelium, where absence of proteases would maintain full-length σ1. Distinct from gut epithelial cells, the levels of SA and JAM-A varies among tumor cells (44–48). Accordingly, we queried whether cleavage of σ1 to σ1N and consequential loss of JAM-binding affected attachment of reovirus to non-epithelial cells. Equivalent particles of PBS- or T.E.E-treated reovirus were exposed to tumorigenic L929 mouse fibroblast cells at 4°C to permit attachment without entry. Cell-associated particles were then enumerated by flow cytometric analysis using anti-reovirus antibodies (Figure 2A). A serial dilution of PBS-treated particles was used for quantitative purposes, and demonstrated a linear relationship between particle number and cell attachment. T.E.E treatment reduced binding of reovirus to L929 cells by ∼80% (Figure 2B).

**Figure 2:**
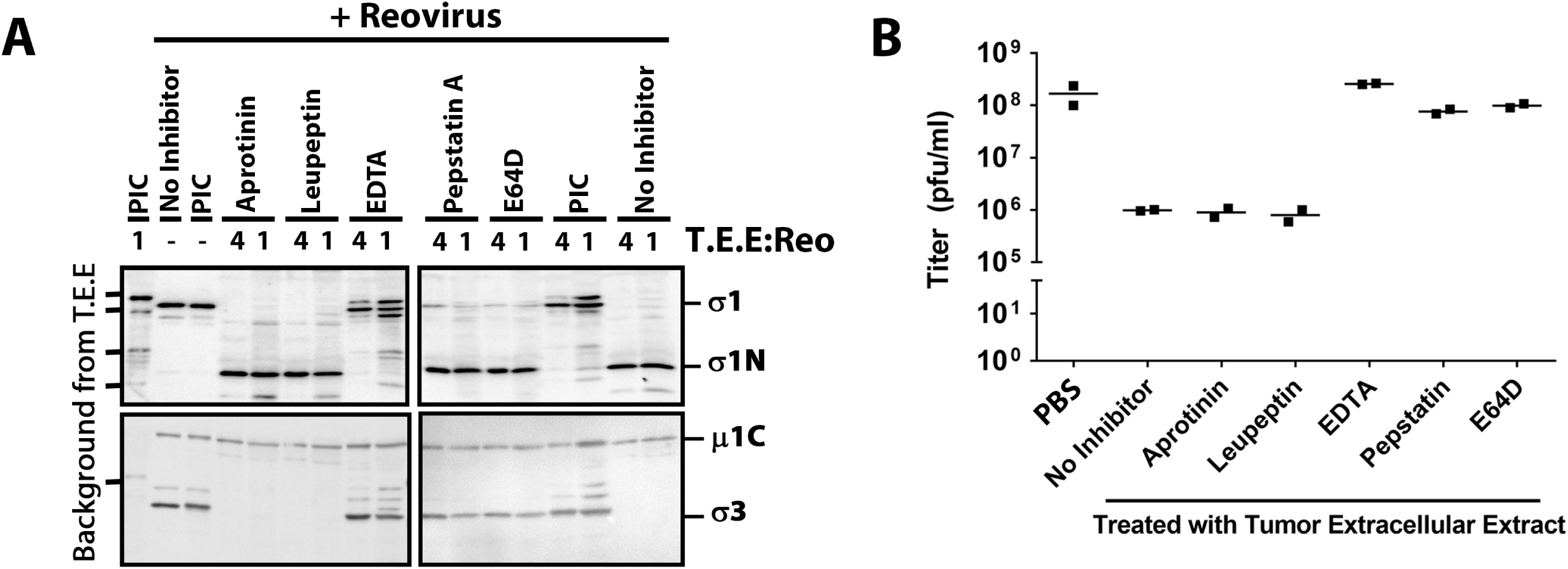
Breast tumor extracellular protease activity reduces reovirus infectivity of L929 cells by 100-fold. **(A)** Attachment to L929 cells assessed by flow cytometry. Cells were exposed to PBS (mock), or reovirus treated with PBS or TE at equivalent does (1x particles) for 1 hour at 4°C, washed, and stained with anti-reovirus and fluorescence-conjugated anti-rabbit antibodies. One-half serial dilutions of PBS-treated reovirus shows linear relationship between mean fluorescence intensity and virus amount and sub-saturation of conditions. **(B)** Same as (A) repeated in 3 independent experiments. To determine relative binding, the MFI of the peak from TE-treated T3D at 1× dilution was extrapolated based on the standard curve generated for serial dilutions of PBS-treated T3D. P<0.0001, student T-test. **(C)** Reovirus was treated with PBS or T.E.E. as in Figure 1 and subjected to plaque titration on L929 cells. The titer (pfu/ml) is presented for three independent mouse tumors, each treated with PBS or T.E.E. five independent times. P<0.0001 between UT and T.E.E. samples, ANOVA statistical analysis and Dunnet’s multiple comparison test.

To determine the impact of T.E.E on infectivity, we then treated reovirus with T.E.E. versus PBS and compared infectious titers (plaque forming units; pfu) on L929 cells commonly used to propagate reovirus. T.E.E. from three independent mouse tumors reduced reovirus titers by ∼100-fold relative to the mock (PBS) treatment (Figure 2C). This finding was striking, as it suggests that tumor proteases may pose a barrier to reovirus oncolytic potency by reducing virus infectivity.

### Mechanistic explanation for loss of infectivity: Truncation of σ1 reduces binding by 100-fold in sialic acid-low cells

Since T.E.E treatment not only caused cleavage of σ1, but also sometimes cause degradation of σ3 and perhaps other effects on protein conformations, it was important to directly determine if σ1 cleavage was necessary and sufficient for decreased reovirus binding. We therefore used reverse genetics to generate reovirus particles with truncated σ1N fibers (Figure 3A, T3D^RG/Δσ1C^) that mimic σ1-cleavage by T.E.E. treatment without effects on σ3. We also generated wild-type reovirus (T3D^RG^) by reverse genetics to control for potential secondary effects of the reverse genetics system. In all experiments, T3D and T3D^RG^ behaved the same, demonstrating that reverse-genetics-generated virions faithfully recapitulate the parental phenotype. Western blot analysis of CsCl-purified virus showed that T3D^RG/Δσ1C^ had truncated σ1N while T3D and T3D^RG^ had full-length σ1 (Figure 3B). All three viruses had equivalent relative levels of µ1C and σ3; therefore, any differences between these viruses can be attributed to σ1 status.

**Figure 3:**
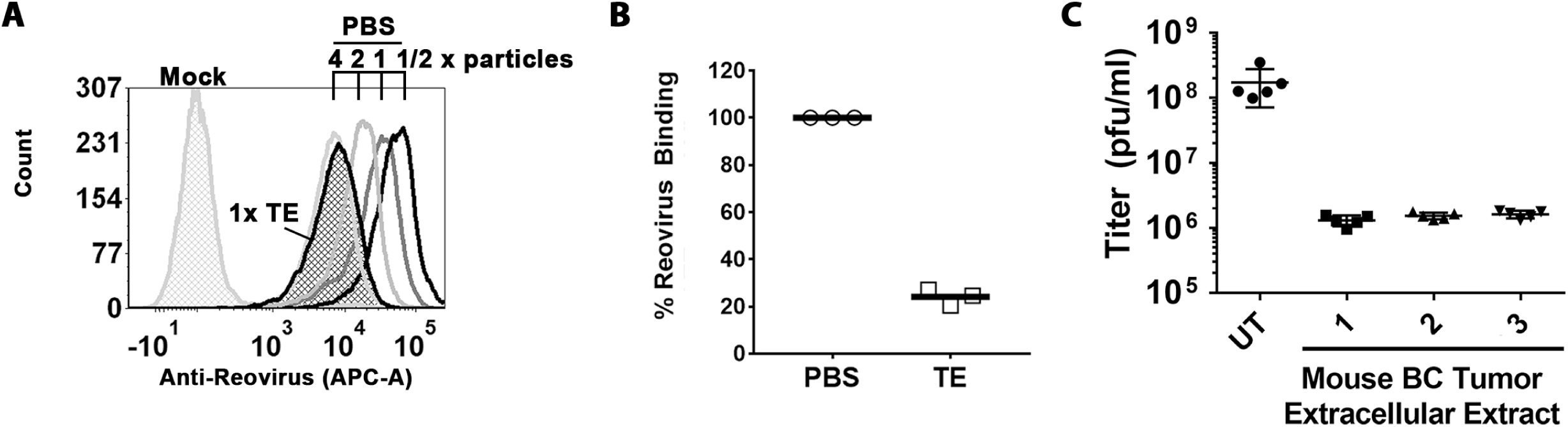
Truncation of σ1 reduces binding of reovirus to sialic acid-low cells by 100-fold. **(A)** Depiction of σ1 truncation in T3D^RG/Δσ1C^. **(B)** Western blot analysis of σ1 and outer capsid proteins of wild type viruses T3D and T3D^RG^ versus mutant reovirus T3D^RG/Δσ1C^, showing that T3D^RG/Δσ1C^ contains truncated σ1N. **(C)** T3D and T3D^RG/Δσ1C^ were treated with PBS or T.E.E similar to Figure 1 and subjected to Western blot analysis for cleaved or truncated σ1 versus other capsid proteins. **(D)** Attachment of T3D and T3D^RG/Δσ1C^ treated with PBS or T.E.E to L929 cells assessed by flow cytometry similar described for Figure 2A. Representative of 3 independent experiments. **(E)** Western blot analysis was used to confirm that at the same dilution, there was equivalent levels of virus proteins and therefore equivalent numbers of T3D, T3^RG^ and T3D^RG/Δσ1C^ purified viruses used for binding assessment. **(F)** Example of flow cytometry to quantify attachment of T3D and T3D^RG/Δσ1C^ to MDA-MB-231 cells. To determine relative binding of T3D^RG/Δσ1C^ to T3D, the MFI of the peak from T3D^RG/Δσ1C^ at 1× dilution produced was extrapolated based on the standard curve generated for T3D. **(G)** Binding assay with T3D^PL^ and T3^RG^ versus T3D^RG/Δσ1C^ reovirus on HeLa, MDA-MB-231, L929, MCF7 cells. Mean fluorescence intensity (MFI) was set to 100% for T3D^RG^ on a given cell line, and remaining virus MFIs presented as a percentage of binding relative to T3D^RG^. Results depict five independent experiments, with 2-way ANOVA statistical analysis and Dunnet’s multiple comparison test. **(H)** Infectivity of T3D, T3^RG^ and T3D^RG/Δσ1C^ in cancer cell lines with different expression levels of JAM and SA, as indicated. Reovirus infection was detected by immunocytochemistry using a polyclonal anti-reovirus serum. Panels shown are representative of two independent experiments **(I)** Sialic acid surface expression detected with fluorophore-labelled SNA for MCF7, MTHJ, TUBO and T47D breast cancer cells. Neurominidase treatment versus untreated cells was used to confirm specificity of SNA staining for sialic acids. **(J)** Mean fluorescence Intensity for SNA staining relative to unstained controls (SNA minus mock), similar to (I), for 2-3 independent experiments. *p<0.05, **p<0.001, ANOVA with Tukey’s multiple comparisons test. **(K)** Mean fluorescence intensity measures the relative particle attachment for T3D versus T3D^RG/Δσ1C^ on MCF7, MTHJ, TUBO and T47D breast cancer cells. Similar calculation as (G) for 2 independent experiments.

If cleavage of σ1 by T.E.E was alone sufficient for reduced binding and infectivity observed in Figure 2, then we expected that T3D^RG/Δσ1C^ should show (1) reduced binding to L929 cells, and (2) should not be affected by T.E.E. treatment. Equivalent particles of T3D^RG/Δσ1C^ versus T3D were therefore assessed for cell-attachment in the presence or absence of T.E.E treatment using flow cytometric analysis (Figure 3C and D). First, T3D^RG/Δσ1C^ exhibited reduced attachment relative to T3D that mimicked levels observed with T3D treated with T.E.E. Second, while T3D binding reduced following T.E.E treatment, attachment of T3D^RG/Δσ1C^ to L929 cells was unaffected. The data implicates cleavage of σ1 as the dominant contributing mechanism for reduced binding of reovirus following T.E.E treatment.

Attachment efficiency was then tested on a panel of cancer cell lines that have differentially express surface JAM and SA; for example, human cervical carcinoma (HeLa) cells and human breast cancer (MDA-MB-231) cells were previously shown to have low JAM- but high SA-surface expression, while L929 cells and the human breast cancer cell line MCF-7 to have low SA expression (48, 49). Figure 3E shows a representative standardization of virus particles and Figure 3E a representative analysis of T3D versus T3D^RG/Δσ1C^ binding to MDA-MB-231. The binding assay was then repeated in five independent experiments, and MFI for a given cell line normalized to T3D^RG^ (Figure 3G). T3D^RG/Δσ1C^ exhibited a statistically significant reduction (∼100-fold) in binding to SA-low L929 and MCF7 cells relative to control viruses, suggesting that loss of the σ1 head domain is sufficient to make reovirus dependent on SA for binding. Moreover, eliminating the JAM-binding σ1 domain in T3D^RG/Δσ1C^ caused reduced infectivity (*i.e.* frequency of cells stained positive for reovirus antigen) towards SA-low L929 and MCF-7 cells, but not towards SA-high HeLa cells (Figure 3H).

The relationship between SA levels and infectivity of reovirus with truncated σ1 was important to validate, as it suggests that not only can the protease status of tumors impact reovirus infectivity, but also the SA status of the tumor cell. We therefore tested two mouse (MTHJ and TUBO) and two human (MCF7 and T47D) breast cancer cell lines for levels of SA versus reovirus attachment. Levels of SA were measured by flow cytometry using fluorescence-labelled Sambucus nigra lectin (SNA). Specificity of fluorescence for SA was confirmed by pre-treatment of cells with neurominidase (Figure 3I). Note that in all cases, neurominidase strongly reduced SNA labelling but did not abolish signal completely, which was expected because neurominidase activity is rarely complete. The four breast cancer cell lines varied dramatically in SA levels, with MCF7 representing minimal SA and T47D maximal SA levels relative to the rest (Figure 3J). Importantly, T3D^RG/Δσ1C^ only exhibited reduced binding relative to T3D when SA levels were low (Figure 3K, MCF7 and MTHJ cells). Together, the findings strongly support that truncation of σ1 reduces attachment of reovirus towards SA-low cells.

### Reovirus-inactivating breast cancer proteases are metalloproteases

Considerable research has demonstrated that tumor environments are rich in proteases of all classes and that proteases can impact the fate of tumor growth, and metastasis (25, 50). To elucidate the class of protease(s) present in T.E.E. that acts on σ1 and σ3, reovirus was treated with T.E.E. in the presence of protease inhibitors that target specific classes of proteases. Aprotinin, leupeptin, pepstatin A, and E64D, were specifically used to inhibit serine, cysteine/serine/threonine, aspartyl and cysteine proteases, respectively (Figure 4A). Since metalloprotease activity depends on metals as cofactors (51, 52), EDTA was used to chelate metal ions and therefore inhibit metalloproteases. The protease inhibitor cocktail (PIC) was able to impair cleavage of σ1 and degradation of σ3, as demonstrated in Figure 1D. Neither aprotinin nor leupeptin were capable of impairing σ1 cleavage and σ3 degradation. Interestingly, Pepstatin A and E64 very minimally impaired σ1 cleavage, but strongly impaired degradation of σ3. EDTA most drastically impaired proteolysis of both σ1 and σ3, suggesting that metal ions are involved, and the dominant protease is a metalloprotease.

**Figure 4:**
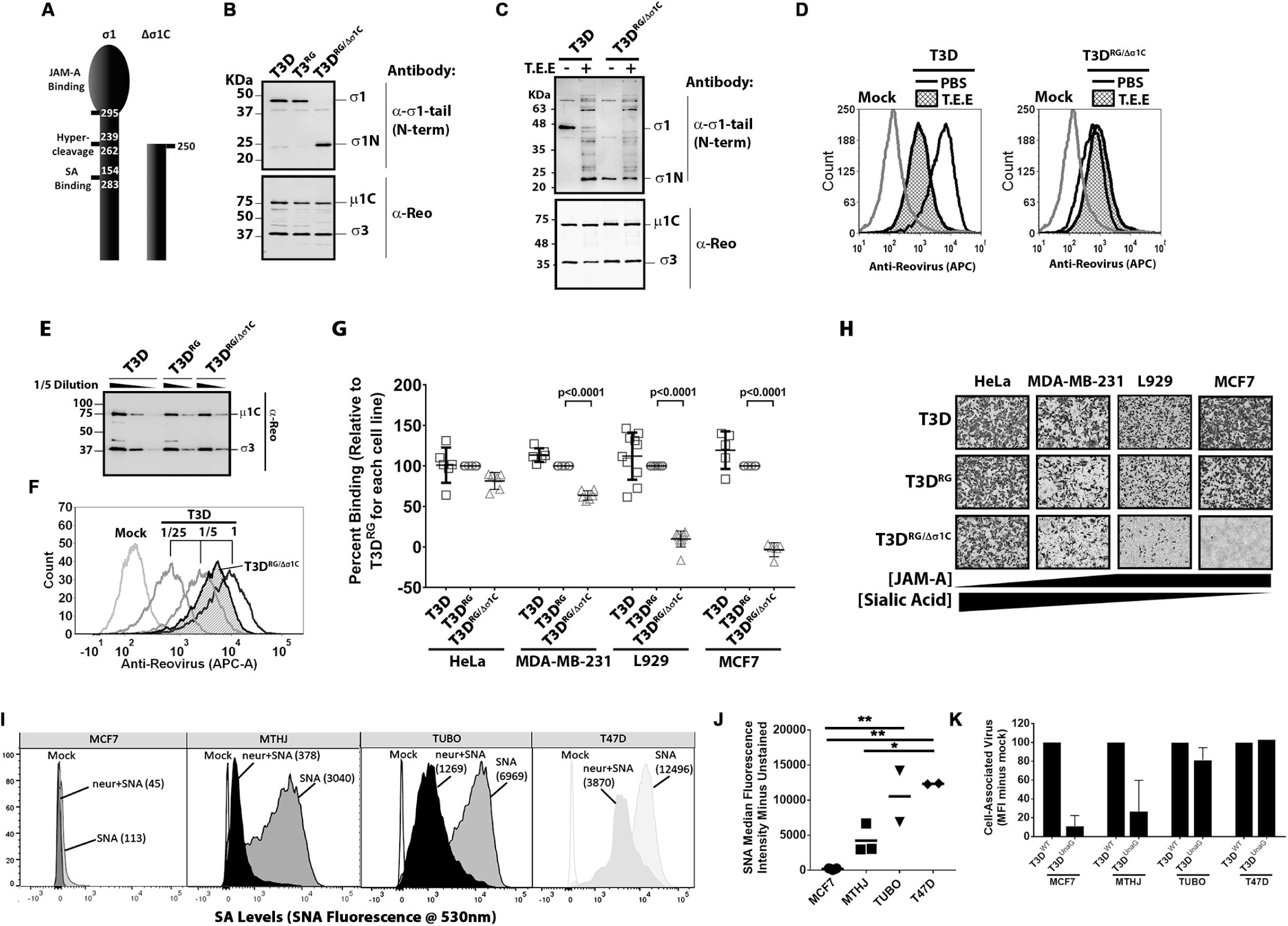
Breast cancer proteases that inactivate reovirus are metalloproteases. **(A, B)** Reovirus was treated with 1× (+) T.E.E. in the presence of various protease inhibitors (as indicated) and analyzed (**A**) by Western blot analysis as in Figure 1 (representative of two independent experiments), or (**B**) by plaque titration on L929 cells as in Figure 2 (two independent experiments).

Next, we examined which protease inhibitors could reverse the loss of reovirus infectivity caused by T.E.E. treatment (Figure 4B). Plaque titration was conducted on L929 cells similar to Figure 2. As seen previously, exposure of reovirus to T.E.E. caused a 100-fold decrease in infectious titers. Neither aprotinin nor leupeptin were capable of rescuing infectivity, as expected given their inability to prevent σ1 cleavage. Pepstatin A and E64 treatments partially rescued infectivity, which correlates with their ability to partially prevent cleavage of σ1. We and others previously demonstrated that although reovirus particles can hold up to 12 σ1 fibers, 3 or more fibers are necessary and sufficient to mediate virus attachment (53, 54). Accordingly, the maintenance of some full-length σ1 during pepstatin A and E64 treatment (Figure 4A) explains why infectious titers remain high (Figure 4B). EDTA was able to fully rescue infectivity, reinforcing that metal ions play a key role in T.E.E.-mediate proteolysis of reovirus σ1 and subsequent loss of infectivity towards SA-low cells.

Together the data suggests that pepstatin A- and E64- sensitive aspartyl and cysteine proteases (respectively) partially contribute to σ1 cleavage, but that metalloproteinase(s) are the main culprits in cleaving σ1 and reducing reovirus infectivity towards SA-low cells.

### Breast cancer cells secrete proteases capable of cleaving σ1

Proteases in the tumor microenvironment can come from multiple sources; from the tumor cells, supporting fibroblasts, or innate and adaptive immune cells (25). To determine if tumor cells directly release a protease that cleaves reovirus σ1 we collected the conditioned media from a panel of cancer cell lines, filtered and concentrated the media 10X using 10 kDa –cutoff centricon columns, and exposed reovirus to these ‘medium extracellular extracts’ (M.E.E.). The rationale for concentrating the media was that cell culturing artificially dilutes extracellular factors by at least 100-fold and therefore can strongly underestimate factor activity. For example, we culture 10^6^ cells in 1ml of media, while T.E.E and I.E.E are collected in 1ml for over 10^8^ cells in tumors (55), which by itself is a dilution of factors that would naturally be more concentrated. Therefore, we initially concentrated M.E.E by 10-fold and since we noted activity, we did not pursue further concentration. Also key to success of this assay was the use of virus production-serum free medium (VP-SFM) to overcome the quenching effects of fetal bovine serum proteins on protease activity. Also, the medium was collected after 3 days of culture when cell death (i.e. floating cells) was minimal. The VP-SFM was largely clear of proteins, as visualized by SDS-PAGE electrophoresis and Coomassie staining (Figure 5A). Conversely, M.E.E from 5 breast (MCF-7, MDA-MB-468, MDA-MB-231, Hs578T and T-47D) and 2 lung (H1299 and A549) cancer cell lines contained a spectrum of secreted proteins (Figure 5A).

**Figure 5.**
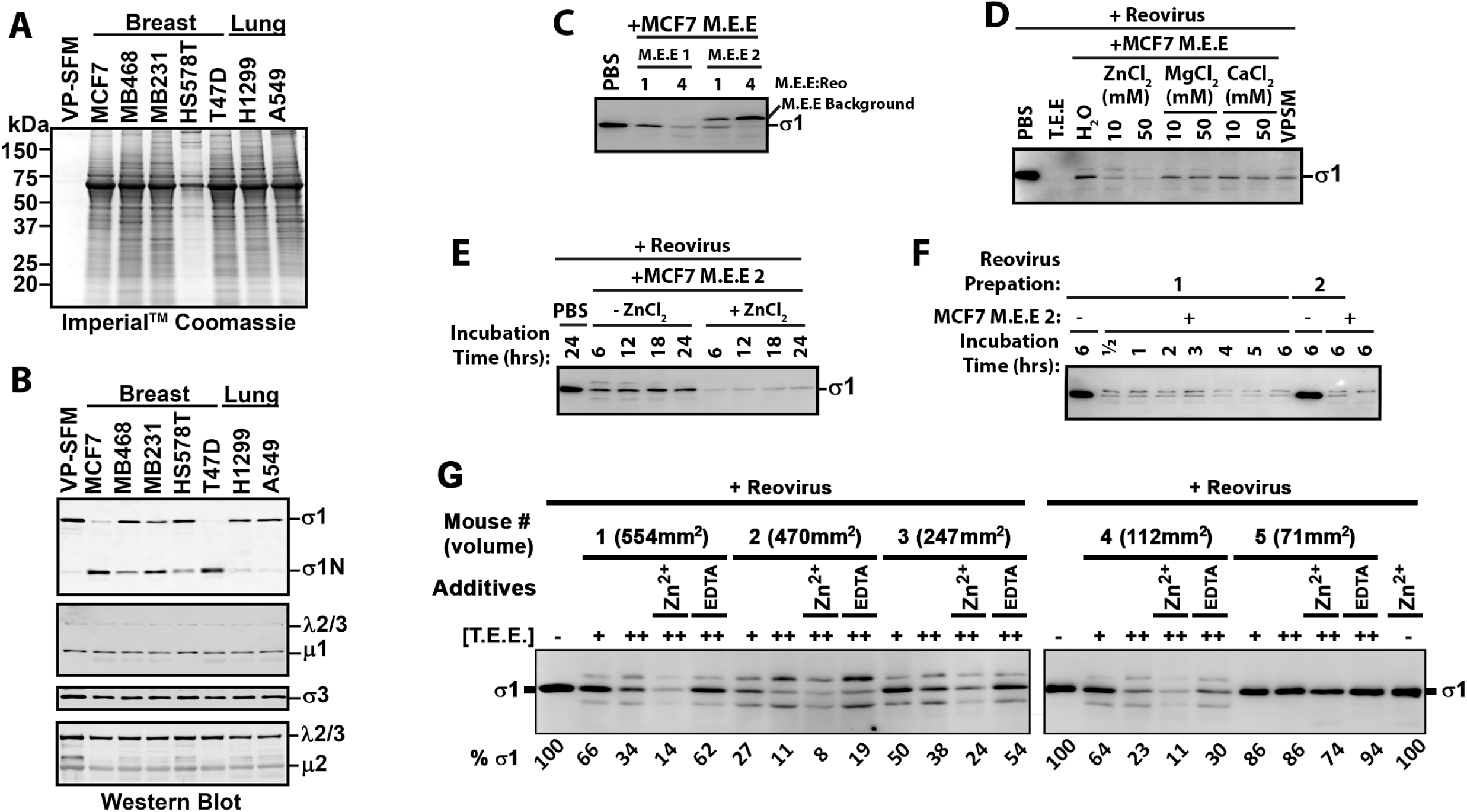
Breast cancer cells secrete proteases capable of cleaving σ1. **(A)** Media extracellular extract (M.E.E.) collected from breast- and lung- cancer cell lines (as indicated) was subjected to SDS-PAGE gel electrophoresis and Coomassie staining to visualize total secreted proteins. **(B)** Reovirus was treated with M.E.E. from indicated cell lines for 24 hours at 37°C and subjected to Western blot analysis as in Figure 1 to monitor the fates of reovirus proteins. Results are representative of 7 independent experiments and 2-4 independent M.E.E. extractions from each cell line. **(C)** Focusing on MCF-7 cells, the reproducibility of M.E.E. activity on reovirus σ1 was evaluated by treatment of reovirus with two additional and independently obtained MCF7 M.E.E. solutions (M.E.E 1 and M.E.E 2), at two different ratios of M.E.E:Reo, and preservation of full length σ1 assessed by western blot analysis. **(D)** To establish which ions contributed to proteolysis of σ1 by MCF-7 M.E.E., reovirus was treated with MCF-7 M.E.E in the absence or presence of ZnCl_2_, MgCl_2_, or CaCl_2_ at indicated concentrations, and preservation of full length σ1 assessed by western blot analysis. **(E)** To determine how rapidly reovirus σ1 is digested by MCF-7 M.E.E, reovirus was treated with MCF-7 M.E.E in the absence or presence of ZnCl_2_ for 6, 12, 18 and 24 hours and preservation of full length σ1 assessed by western blot analysis. **(F)** To ensure that susceptibility to MCF-7-mediated digestion of σ1 was independent of the reovirus preparation, two independently-purified reovirus stocks were treated with MCF-7 M.E.E. or PBS for 0.5 to 6 hours (as indicated), and preservation of full length σ1 assessed by western blot analysis. **(G)** MCF7 tumors implanted into NSG mice were excised after 45 days of *in vivo* tumor growth and T.E.E. was prepared. Reovirus was treated with 1x and 2x T.E.E., or 2x T.E.E in presence of EDTA or ZnCl_2_ and subjected to Western blot analysis. Densiometric analysis of full-length σ1 is provided below the blot relative to PBS treated reovirus.

When treated with M.E.E. from all 5 breast cancer cell lines, reovirus σ1 underwent cleavage to σ1N (Figure 5B) but limited change was observed for µ1C. MCF7 M.E.E. collections from two independent cultures showed cleavage of σ1 to different extents, suggesting that the concentration of proteases varies between cultures (Figure 5C). Overall, breast cancer cell M.E.E. recapitulated the effects of T.E.E., indicating that tumor cells alone can secrete proteases that target reovirus σ1. Our findings of course do not eliminate the possibility that supporting fibroblasts, or innate and adaptive immune cells in tumor microenvironments also contribute additional reovirus-inactivating proteases. It was interesting that neither lung cancer cell M.E.E. cleaved reovirus σ1. Although tempting to speculate that tumor cell type (*e.g*., breast versus lung) may influence differential ability to inactivate reovirus through secreted proteinases, a larger panel of lung cancer cells would be essential to make such interpretation.

Reovirus was also treated with MCF-7 M.E.E in the absence or presence of ZnCl_2_, MgCl_2_, or CaCl_2_ to determine which metal ion promoted σ1 cleavage. With this stock of MCF-7 M.E.E., σ1 was completely degraded rather than producing a stable σ1N fragment, as was described with high T.E.E. concentrations (Figure 1B), so we therefore focused on the loss of full-length σ1 as our readout. Importantly, degradation of σ1 was reproducibly increased in the presence of Zn^2+^, implicating a Zn-dependent MP in reovirus processing (Figure 5D). A time-course analysis showed that σ1 proteolysis occurred rapidly by M.E.E. (Figure 5E), reaching maximum loss of full-length σ1 by 6 hours of treatment, with Zn^2+^ again promoting σ1 processing. A more detailed time course analysis shows maximal proteolysis achieved already by 30-minute exposure of two independent preparations of reovirus to MCF-7 M.E.E (Figure 5F). Together, the data suggest that breast cancer cells can themselves secrete Zn-dependent metalloproteases that cleave σ1 with rapid kinetics.

Finally, we determined if xenograft tumors of MCF7 cells in severely immune-compromised NOD scid gamma mice (lacking mature T cells, B cells and natural killer cells) produced a tumor microenvironment capable of cleaving reovirus σ1 (Figure 5G). At 45 days post-implantation, excised tumors were rinsed twice in PBS, cut into 4 pieces, and incubated at 4°C in 1 ml (total) PBS for 2 hours to diffuse extracellular content. These tumor extracellular extracts (T.E.E.) were clarified by centrifugation and 0.45um filtration. Reovirus was then exposed to the T.E.E.s and loss of full-length σ1 was monitored by Western blot analysis (Figure 5G). Degradation of σ1 was T.E.E. dose-dependent and varied from ∼10-90% depending on the tumor. Four of the five T.E.E.s showed >60% cleavage. Moreover, there seemed to be a relationship between the size of tumor and cleavage efficiency, although many more samples would be required to strengthen the correlation. Since cleavage was increased in the presence of Zn^2+^ and decreased in the presence of EDTA, a Zn-dependent metalloprotease is likely functioning in these tumors. However, since EDTA treatment did not completely prevent cleavage, it is also possible that additional ion-independent σ1-degrading proteases were active in MCF7 tumors. These experiments confirm the presence of σ1-cleaving metalloproteases in tumors *in vivo*, and in absence of T-, B-, or NK- cell sources.

### Mutation of the σ1 neck domain overcomes reovirus cleavage by breast cancer metalloprotease

Given that proteolysis of reovirus σ1 by Zn-dependent metalloproteases secreted by human breast cancer cells and present in mouse breast cancer tumors diminished reovirus binding and infection towards SA-low tumor cells, we next sought to modify reovirus σ1 to overcome cleavage. Proteolysis of reovirus by intestinal proteases chymotrypsin and trypsin was previously shown to occur in the flexible protease-hypersensitive “neck” region (residues 219-264) between the tail and the head domains of σ1 (56, 57) (Figure 6A). We reasoned that if tumor-associated metalloproteases also cleaved in the σ1 neck domain, then the σ1N fragments generated by T.E.E., I.E.E., trypsin, and chymotrypsin should share a similar molecular weight. Indeed, the tail (σ1N) and head (σ1C) fragments detected by Western blot analysis with σ1N- and σ1C-specific antibodies (respectively) were similar when reovirus was treated with T.E.E., I.E.E., trypsin, or chymotrypsin (Figure 6B). The shorter fragments of σ1 under some conditions of I.E.E and chymotrypsin treatment is likely to reflect further cleavage at the extreme N-terminus as described previously (58, 59). Importantly, since chymotrypsin and trypsin cleavage sites are six amino acids apart but the size of their σ1 cleavage fragments are not resolved by our assay, we can infer that the tumor-associated MP cleaves in the same general vicinity as the gut proteases.

**Figure 6:**
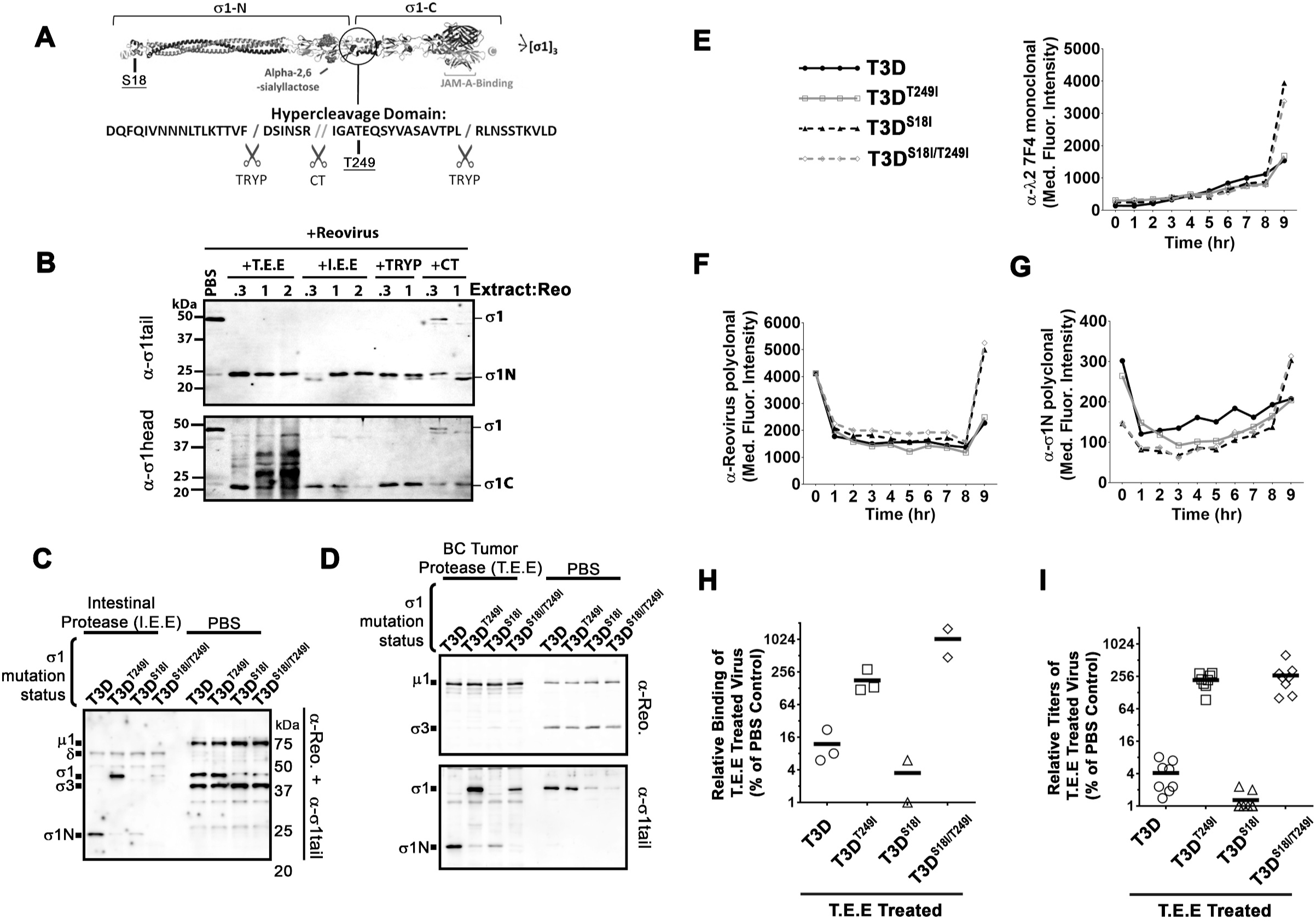
Mutation in the σ1 neck domain can overcome σ1 proteolysis by breast cancer metalloprotease. **(A)** Diagrammatic depiction of **σ**1 with the protease-hypersensitive neck domain. **(B)** Reovirus was treated with either T.E.E., I.E.E., chymotrypsin (CT) or trypsin (TRYP) for 24 hours at 37°C and subjected to Western blot analysis with tail- (top) or head- (bottom) specific antibodies. **C - D.** CsCl-purified T3D, T3D^T249I^, T3D^S18I^ or T3D^S18I/T249I^ were treated with **(C)** PBS, I.E.E or **(D)** T.E.E. for 24 hours at 37°C. Western blot analysis with both polyclonal anti-reovirus antibodies and σ1N-specific antibodies demonstrate the levels of full-length σ1 and σ1N. (**E-G)** Reovirus infection dynamics of T3D, T3D^T249I^, T3D^S18I^ or T3D^S18I/T249I^ viruses. Flow cytometry was used to evaluate expression of reovirus proteins: **(E)** λ2, **(F)** σ3, **(G)** σ1N, from 0 to 8 hours post infection. **(H)** Binding assay as in Figure 3 with reovirus mutants: T3D, T3D^T249I^, T3D^S18I^ or T3D^S18I/T249I^ treated with T.E.E. on L929 cells. **(I)** Plaque titration of T.E.E. treated reovirus mutants (T3D, T3D^T249I^, T3D^S18I^ or T3D^S18I/T249I^) as in Figure 2.

It was previously observed that a change from threonine to isoleucine at position 249 of σ1 can prevent cleavage by both chymotrypsin and trypsin despite their different cleavage locations in the neck domain (56). These findings suggested that the T249I modification eliminated cleavage susceptibility by altering the secondary structure of the neck domain, thereby altering the exposure of the hyper-cleavage domain. Accordingly, we tested if mutation of T249 to isoleucine could also prevent cleavage by tumor-associated metalloproteases. Using reverse genetics, we introduced the T249I mutation into T3D and assessed the fate of σ1^T249I^ after treatment with I.E.E. (Figure 6C) or T.E.E. (Figure 6D). The T249I mutation successfully impeded cleavage of σ1 by both I.E.E. and T.E.E.

Our previous studies showed that a mutation in the domain that anchors σ1 in virions, σ1-S18I, reduces the number of σ1 fibers per reovirus particle to ∼4 (instead of 12 on wild-type T3D). We and others further showed that 3 σ1-trimers were sufficient to allow maximal binding to L929 and other tumorigenic cells. Moreover, having 4-7 (but fewer than 12) σ1 trimers promotes uncoating of σ1 during virus entry into tumor cells, and thereby increases reovirus oncolysis *in vitro* and *in vivo* (53, 60). Having now learned that σ1 undergoes cleavage by breast tumor-associated metalloproteases, we reasoned that having fewer σ1 fibers would make T3D^S18I^ hypersensitive to tumor-associated protease inactivation; in other words, that maintaining full-length σ1 would become less-likely if there were fewer σ1 fibers to begin with. We therefore also incorporated the T249I mutation into T3D^S18I^ to generate a double-mutant T3D^S18I/T249I^. As expected, both T3D^S18I^ and T3D^S18I/T249I^ showed lower σ1 levels relative to T3D or T3D^T249I^ (Figure 6C and 6D). Importantly however, while σ1 of T3D^S18I^ was cleaved by I.E.E. and T.E.E., the σ1 T3D^S18I/T249I^ was refractory to cleavage by either extracellular extract. In summary, localizing the σ1 cleavage site to the neck domain, and subsequently introducing a T249I mutation, allowed us to successfully generate T3D and T3D^S18I^ variants that withstand proteolysis by breast tumor-associated metalloproteases.

Chappell et al (1998) previously found that clinical reovirus strains exhibit either a threonine or isoleucine at position 249 of σ1 (56), but it was unknown whether a T249I mutation would have undesired determinantal effects on virus entry, endocytosis, or uncoating. To test this, L929 cells were exposed to equivalent particle doses of T3D, T3D^T249I^, T3D^S18I^ or T3D^S18I/T249I^ at 4°C, washed extensively, then incubated at 37°C for 0-9 hours. At every hour, cells were fixed, stained for specific reovirus proteins, then analyzed by flow cytometry to follow the fate of input reovirus particles versus *de novo* reovirus protein expression. First, flow cytometry with λ2-specific antibodies, which cannot detect input virions (*i.e.,* λ2 epitopes are hidden in the virion) but can detect *de novo* λ2 protein expression, demonstrated new virus protein expression at 8 hours post-infection (hpi). Importantly, T3D and T3D^T249I^ demonstrated similar *de novo* protein synthesis levels, suggesting similar kinetics of infection (Figure 6E). T3D^S18I^ was similar to T3D^S18I/T249I^ with respect to *de novo* λ2 expression. As expected from previous studies showing that S18I increases reovirus infectivity (53, 60), both T3D^S18I^ and T3D^S18I/T249I^ exhibited ∼3-fold more *de novo* λ2 expression relative to T3D and T3D^T249I^.

Next, polyclonal anti-reovirus antibodies and monoclonal antibodies towards σ3 that detect both input virions and *de-novo* virus protein synthesis, confirmed equivalent input levels for all four viruses, yet increased infectivity (or rate of infectivity) of variants containing the previously-characterized S18I mutation (Figure 6F). Again, it is important to note that the T249I mutation did not impact the efficiency of establishing infection. Finally, antibodies directed to the tail domain of σ1 confirmed that input virions containing the S18I mutation contained ∼3-fold less σ1 but produced more *de-novo* proteins (Figure 6G). Altogether these results indicate that the T249I mutation does not negatively affect T3D reovirus infection whether in the context of wild-type T3D or the more oncolytic T3D^S18I^ variant.

Since cleavage of σ1 by T.E.E. reduced attachment to SA-low cells and inhibited infectivity (Figure 2 and 3), we evaluated if the T249I mutation that prevents σ1 cleavage can facilitate reovirus infectivity in the presence of MPs. Accordingly, L929 cells were exposed to equivalent doses of T3D, T3D^T249I^, T3D^S18I^ or T3D^S18I/T249I^ that had been pretreated with T.E.E., then binding was evaluated by flow cytometry (Figure 6H), and infectivity evaluated by plaque assays (Figure 6I). T3D^T249I^ bound to L929 cells 16X more than T3D (Figure 6H), indicating that it was resistant to protease cleavage. This increase in binding correlated with 16X higher virus production than T3D (Figure 6I). T3D^S18I^ showed lower binding and virus production than T3D probably because this mutant has reduced σ1 levels and is therefore hypersensitive to σ1 cleavage. Importantly, the dysfunction in binding and virus production was overcome by combining with the T249I mutation in T3D^S18I/T249I^. These results suggest that incorporating the T249I mutation into T3D generates a virus capable of resisting T.E.E.

### T3D^T249I^ and T3D^S18I/T249I^ do not have added toxicity relative to T3D in *vivo*

Clinical trials using T3D as a monotherapy in several cancers have shown that T3D is a safe therapy but would benefit from enhanced efficacy (61). Little is known about the effects of the tumor environment on reovirus oncolytic performance. Having found that T.E.E. cleaves σ1 and reduces infectivity towards tumor cells with low sialic acid levels (Figure 3), and having developed σ1- “uncleavable” variants of T3D (T3D^T249I^ and T3D^S18I/T249I^) (Figure 6), suggests that uncleavable reovirus may warrant consideration for testing as oncolytic agents. The first priority, however, was to determine if these mutants were safe. Previous studies found that JAM-A on endothelial cells contributed to bloodstream dissemination of serotype 1 reovirus (62), so it seemed essential to test if T3D^T249I^ and T3D^S18I/T249I^, by virtue of retaining JAM-A binding, pose any toxicity / safety concerns relative to T3D.

While reovirus is restricted to tumors and safety has been demonstrated in immunocompetent mice and in humans in clinical trials, reovirus shows visible signs of toxicity in severely immune-compromised NOD scid gamma (NSG) mice. NGS mice lack mature T cells, B cells and natural killer (NK) cells. In the absence of immune restrictions, T3D reovirus can disseminate from tumors to heart and circulatory systems, impair circulation, and cause black-foot syndrome (63). We therefore tested the toxicity of T3D strains when injected into MCF7 tumor xenografts in the background of NSG mice. Specifically, MCF7 cells were implanted into the mammary fat pads of NSG mice. When tumors became palpable, five mice were injected intratumorally with either PBS, or 10^6^ plaque-forming-units (PFUs) of T3D, T3D^T249I^, T3D^S18I^ or T3D^S18I/T249I^. The dose was chosen based on our preliminary dose escalation experiments with T3D, which defined 10^6^ as an optimal dose to see toxicity within guidelines of strict animal care protocols (data not shown). Reovirus injections were repeated for a total of three times over a one-week period. Mice were monitored and euthanized based on humane endpoints (first sign of black tail/black foot, or over 15% weight loss). The onset of symptoms of toxicity for the different treatment groups is shown in Figure 7A. Although time to symptoms varied between 25 and 32 days after the first virus inoculation, there were no significant differences among the virus treatments with respect to appearance of symptoms. PBS-treated animals did not exhibit symptoms of toxicity and were euthanized at 45 days. Reovirus titers in the hearts were not significantly different between groups (Figure 7B). Therefore, in the NSG model of reovirus toxicity in absence of sufficient immune restriction, the σ1-uncleavable T3D variants did not pose additional toxicity or safety concerns relative to wild-type T3D.

**Figure 7:**
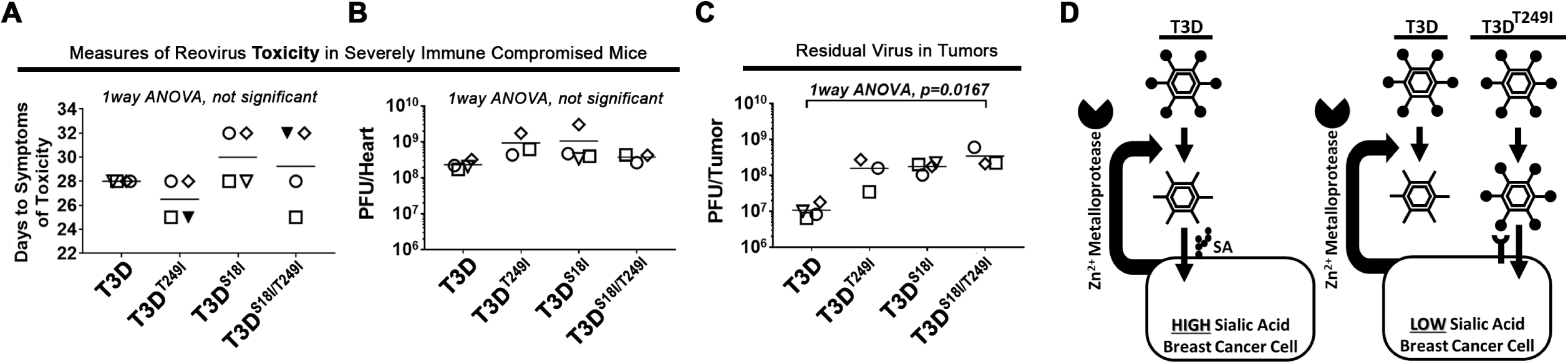
Uncleavable reovirus does not increase toxicity in NSG mice when implanted intra-tumorally. **(A)** Toxicity of T3D, T3D^T249I^, T3D^S18I^ or T3D^S18I/T249I^ was assessed in severely immune compromised (NSG) mice. When tumors became palpable, five mice were injected intratumorally with 10^6^ plaque-forming-units (PFUs) of virus. Mice were euthanized when reovirus toxicity was observed (days to symptoms of toxicity), specifically when they showed first signs of black-foot (and/or tail or ears) indicating circulation deficiencies, or lost more than 15% of body weight. We were unable to obtain tissues from the 2 mice indicated by solid-black triangles. In B-D, individual mice within each group have a unique symbol. PBS control injected mice were symptom free at day 45 when experiment was terminated. **(B)** Whole hearts were homogenized and subjected to plaque titration. Titers of reovirus in the heart provide a secondary measure of reovirus-induced toxicity. **(C)** Reovirus titers in whole tumors as in B. **(D) Final model:** Some breast cancer cells secrete zinc-dependent metalloproteases that cleave σ1, removing the JAM-A-binding head domain, and thereby making reovirus dependent on the presence of sialic acids for cell attachment. In breast cancer cells with low levels of sialic acid, the cleavage of σ1 strongly suppresses virus attachment and infection. Introduction of a T249I mutation into σ1 overcame the inactivation of reovirus by tumor-associated proteases, permitting efficient binding and entry.

The onset of black-foot syndrome necessitates euthanasia, which prevented assessment of virus-mediated increase in long-term survival of MCF7 tumor-bearing mice. However, since tumors were available to us upon euthanasia, we did assess the relative infectious virus titers of σ1-uncleavable T3D variants versus wild-type T3D in tumors. The mean reovirus titers in homogenized tumors were 1.0×10^7^, 1.6×10^8^, 1.8×10^8^, and 3.5×10^8^ PFUs for T3D, T3D^T249I^, T3D^S18I^ and T3D^S18I/T249I^ respectively (Figure 7C). While the trend suggested increasing titers for progressive addition of T249I and S18I mutations, statistical significance was only achieved for T3D^S18I/T249I^ versus T3D, which showed a 30-fold difference (p=0.0167). As explored further in the discussion, these studies were designed to monitor safety, and therefore distinct *in vivo* experimental designs beyond the scope of this study would be necessary in future to compare oncolytic properties of these T3D variants. The current animal experiment supports that T3D^S18I^ and T3D^S18I/T249I^ do not exhibit gain of function with respect to virulence when introduced intra-tumorally.

## DISCUSSION

While reovirus undergoes clinical testing as a cancer therapy, it is important to gain fundemintal understanding of factors in the virus and tumor environments that could alter reovirus behavior in tumors. Such factors might contribute to heterogenous response between patients, and/or could help guide strategies to improve the potency of reovirus and other virus cancer therapeutics. Being a proteinaceous entity, we wondered if reovirus was processed by proteases that exist in breast tumor microenvironments. Moreover, since reovirus naturally exploits gut proteases chymotrypsin and trypsin, or lysosomal cathepsins, to partially uncoat into membrane-penetrating ISVPs, we wondered what effect tumor proteases would have on reovirus infectivity towards tumor cells. Our studies show, for the first time, that proteases in breast tumor microenvironments have potential to restrict reovirus infectivity (Figure 7D). Specifically, proteases from *in vivo* breast tumors (T.E.E.) and from breast cancer cell lines (M.E.E) fail to generate ISVPs. Instead, the T.E.E.- and M.E.E-associated proteases cleave the reovirus cell attachment protein σ1 into two portions, removing the C-terminal domain that can bind to JAM-A and other host proteins, while generating reovirus particles that retain only the sialic-acid binding tail domain. In other words, the tumor proteases make reovirus exclusively dependent on sialic acids for binding. Accordingly, T.E.E.- and M.E.E- treated reovirus particles have ∼100-fold less attachment and infectivity towards tumor cells with reduced sialic acid levels. Our analysis further demonstrated that reovirus-inactivating proteases are commonly secreted by breast cancer cell lines, and that the proteases are of the zinc-dependent metalloprotease (MP) family. Moreover, the inactivation of reovirus by breast cancer-associated MPs could be fully overcome by introducing a mutation (T249I) into σ1. Reovirus with the σ1^T249I^ mutation did not exhibit added toxicity in immunocompromised NSG mice, and can therefore proceed to testing for oncolytic activity in immunocompetent breast cancer models. Conceptually, our findings extend beyond reovirus, suggesting that tumor-associated proteases warrant consideration when using proteinaceous therapies in general.

Proteases are prevalent components of the tumor microenvironment, participating in extracellular matrix degradation, tumor cell invasiveness, as well as altering activities of important signaling molecules (64). Many cells contribute to the pool of proteases in tumors, including the tumor cells themselves, mesenchymal stem cells, tumor-supporting fibroblasts and endothelial cells, and both innate and adaptive immune cells. In our studies, we discovered that breast cancer cells can secrete reovirus-inactivating proteases into the medium; but our findings do not preclude the possibility that other cells contribute additional proteases that act on reovirus *in vivo*. It would be interesting to expose reovirus to medium from tumor-supporting cells and immune cells, to determine whether the composition of a specific tumor could predict reovirus processing/infectivity at the tumor site. Nevertheless, the extracellular extracts from two distinct *in vivo* breast tumor models (*i.e.* MCF7-derived tumors and polyoma virus middle T-antigen-derived tumors) produced the same outcome as media derived from breast cancer cells: an absence of the beneficial processing of reovirus to ISVPs, but rather inactivation of reovirus infectivity towards sialic-acid-low tumor cells by cleavage of σ1. Our preliminary analysis showed that unlike breast cancer cells, two lung cancer cell lines failed to secrete proteases that process reovirus. An analysis of a larger panel of lung and other cancer cell lines would be necessary, however, to determine if the inactivation of reovirus is breast-cancer-specific, or can extend to other cancers.

Human proteases are categorized into aspartic, cysteine, metallo, serine, or threonine protease classes, on the basis of the catalytic amino acid (or ion) in the active site. To distinguish which class(es) of proteases cleave σ1, reovirus processing by breast cancer proteases was tested in the presence of pharmacological inhibitors or divalent ions. The experiments indicate that the primary protease activity that cleaves σ1 is a zinc-dependent metalloprotease. As there are 100s of possible candidates, and specific pharmacological inhibitors are only available for a select few, it requires a proteomics approach in the future to identify the precise protease that cleaves σ1. The identity of the protease cannot be determined from the sequence of σ1 due to lack of defined linear consensus sequences for most candidates and yet unclear preferences for specific substrate secondary and tertiary structures. Moreover, it is important to note that our experiments also implicated aspartyl and cysteine proteases as having partial effects on σ1 cleavage and reovirus infectivity (Figure 4). A remarkable feature of the tumor proteolytic network is the large extent to which proteases cleave other proteases and thereby affect their activities. For example, many metalloproteases are themselves substrates of aspartic, cysteine, and serine proteases (64). It is therefore possible that aspartyl and cysteine proteases cleave σ1 directly, or, that they contribute to activation of the key metalloprotease. If the precise protease(s) that cleave σ1 are identified, it would be interesting to test if they serve as predictive markers for the extent of reovirus amplification in breast tumors. Given the complexity of the proteolytic network, we instead focused our attention on characterizing the effects of the protease on reovirus, and modifying reovirus to withstand the protease(s) irrespective of their identity.

Another interesting detail from our studies is that breast cancer MPs make reovirus dependent on sialic acids (SA), and therefore reduce reovirus binding and infection specifically in SA-low breast cancer cells such as MCF-7. SAs are frequently modified in BC (65), for example with overexpressed complex β-1,6-branched glycans (66), incomplete glycan structures (67), or non-mammalian ‘xenoglycans’ (68). Hypersialylation is frequently seen in highly-invasive breast cancers, which increases shedding of SAs that could occlude SA binding sites on the virion. To fully appreciate the effects of σ1 cleavage on reovirus oncolysis in breast cancer, it seems necessary to better understand which sialic acid modifications impact reovirus infection, and whether shed SAs compete for reovirus attachment to cells. The nature of SAs in specific cancers could then potentially serve as predictive markers for reovirus amplification. In any case, by overcoming σ1 cleavage with a T249I mutation, the retention of JAM-A binding capacity may overcome barriers posed by SA insufficiency or soluble SA competition.

As for application of knowledge towards improving reovirus oncolysis, our findings suggest a compendium of possibilities to be tested in the future. First, the T3D^T249I^ and T3D^S18I/T249I^ variants need to be thoroughly compared to T3D parental virus in various *in vivo* animal models, to appreciate the extent to which cleavage of σ1 by tumor proteases affects oncolysis. Next, we propose that modification of T3D to transition to ISVPs in the context of well-characterized tumor proteases could be dually beneficial. Others have previously proposed using ISVPs (rather than whole reovirus particles) for cancer therapy (69); but such a procedure would only benefit infection for the incoming inoculum. Having reovirus consistently transition to ISVPs in tumor environments should produce a benefit at every round of virus amplification. Another idea is to generate reovirus particles that encode natural metalloprotease inhibitors, such as the small tissue inhibitors of metalloproteinase (TIMPs). Metalloproteases have served as possible cancer therapeutic targets, but unfortunately they also play important roles in healthy tissues and therefore global inhibition can be toxic. Perhaps a virus that directly delivers TIMPs to tumor microenvironments could serve two functions; inhibit inactivation of the virus, and prevent pro-tumor activities of MPs. Finally, our findings could extend beyond reovirus, as there are many oncolytic viruses and other proteinaceous treatments such as antibodies undergoing clinical testing and use for cancer therapy. An expanded understanding of the effect(s) of tumor proteases on proteinaceous treatments could help develop second generation therapies with augmented potencies.

Finally, beyond implications on reovirus oncolysis, our finding that proteases in tumor versus intestinal microenvironments exhibit different on reovirus demonstrates that virus-host interactions are not just limited to the target infected cells, but to the entire environment of a niche. In other words, if we better understand extracellular host factors that impact viruses, we may better appreciate why viruses thrive in specific host niches. In addition to proteases, extracellular factors such as the microbiota and immunological host factors can impact viruses. It will be interesting to know whether additional extracellular factors such as the extracellular matrix, secreted lipids, carbohydrates, enzymes or other soluble factors impact the dynamics of viruses and therefore their oncolytic or pathogenic potential.

## Materials and Methods

### Reovirus production

Mammalian Orthoreovirus Serotype 3 Dearing (T3D) was propagated in L929 spinner cultures, extracted, and CsCl purified as previously described (70). Virus titers were determined by plaque assay on L929 cells as previously described (71). The titer of the stock virus used in most experiments was 1.01×10^10^ PFU/mL.

### Cell culture

L929 cells were cultured in minimal essential medium (MEM) (Sigma) supplemented with 10% FBS (Sigma), 1× sodium pyruvate (Sigma), non-essential amino acids (NEAA) (Sigma), and 1× Antibiotic-Antimycotic (Gibco, catalogue #15240062). HeLa cells were cultured in Dulbecco’s minimal essential medium (DMEM) and 1× Antibiotic-Antimycotic. MCF-7, T47D, MB-468, MB-231, H1299, A549 cells were cultured in Roswell Park Memorial Institute (RPMI) 1640 medium (supplemented as described for MEM). L929, Hela, and A549 cells were laboratory stocks obtained from Dr. Patrick Lee’s laboratory (Dalhousie University), while remaining cells were purchased from the American Type Culture Collection; all cells were confirmed mycoplasma free by Hoechst DNA Staining. For experiments, cells were cultured in 1ml per well of a 12 well plate, and media was scaled according to the surface area (for example, in a well of a 6 well plate, we used 2ml of media).

### Preparation of protease rich extracts

To mimic the tumor and intestinal environments reovirus would encounter, we obtained extracellular extracts from both environments. Tumor extracellular extract (T.E.E.) was generated by establishing breast tumors in mice from polyoma virus middle T-antigen derived tumor cells (72). Tumors were explanted and incubated in 1ml phosphate buffered saline at 4°C for 2 hours. The PBS solution was then centrifuged at 10,000 × g and filtered (0.45µm) to reduce cellular debris. I.E.E was made by flushing entire murine intestines with 1ml PBS followed by filtering (0.45µm) to remove debris. Polyoma virus middle T-antigen Tumor sizes ranged from 700-2000mm^2^. The same method was used to extract T.E.E. from MCF7 tumors in Figure 5G, except tumors were smaller and therefore diffused into 0.5ml of PBS (tumor sizes indicated in Figure). To determine whether cancer cells alone are capable of secreting proteases that inactivate reovirus, protease-rich extracellular extract was obtained from a variety of breast and lung cancer cell lines (Breast- MCF-7, T47D, MB-468, MB-231; Lung-H1299, A549) for 24 hours at 37°C (M.E.E; media extracellular extract) using the following protocol: First, cells were cultured to 80% confluent as described above. After an overnight incubation, the supplemented RPMI medium was replaced with protein-free VP-SFM (Virus Production Serum Free Media) supplemented with L-glutamine, and the cells were then cultured for 3 days; health of cells with limited cell detachment was confirmed daily by microscopy. The medium was then collected and centrifuged at 1000 x g for 7 minutes to remove any floating cells and debris. The supernatants were passed through a 0.45μm low protein binding syringe filter. Then, the filtrate was centrifuged in a 5K NMWL (nominal molecular weight limit) centrifugal filter (Millipore UFV4BCC00) at 3000RPM for 5-10 minutes, until volume reached 1/10^th^ initial volume. For use as a negative control, a plate containing supplemented RPMI medium without any cells was subjected to the same procedure. To avoid enzymatic inactivation as a result of freeze-thaw cycles, T.E.E., M.E.E., and I.E.E. were aliquoted before flash freezing in liquid N_2_ and stored at -80°C. A fresh aliquot was used for each experiment

### *In-vitro* proteolysis assay with T.E.E, M.E.E, I.E.E, chymotrypsin or trypsin

Reovirus was incubated at 37°C for 24 hours (unless indicated otherwise) with T.E.E, M.E.E. or I.E.E in the presence or absence of different inhibitors. Volumes of these protease rich extracts varied in each experiment as indicated in figures, but ranged from 1-15µl of extract at original concentration, and 1.08×10^10^ to 5.40×10^10^ viral particles (1-5µl of reovirus, at a concentration of 1.08×10^13^ particles/mL, as calculated based on the equivalency that a spectrophotometric optical density of 260 nm (OD_260_) of 5.42= 1.13×10^13^ particles). The proteolysis assay was conducted in PCR tubes and incubated using a Bio-rad T100 Thermal Cycler (with heated lid). For the timepoint digests Bio-rad S1000 and C1000 thermal cyclers were also used. Reactions were incubated at 37°C for 24 hours, or for the time lengths indicated during timecourse analyses. The inhibitors used were EDTA (ethylenediaminetetraacetic acid at 36.25μM), EDTA-free protease inhibitor cocktail (PIC, 1x final concentration; Sigma-Aldrich: Cat# P8340), and protease inhibitor cocktail with EDTA (CPIC, 1x final concentration; Roche: Cat# 11 697 498 001). Protease-class inhibitors were added to reactions as follows: leupeptin (100µM final concentration, Sigma), aprotinin (200nM final concentration, Sigma), and pepstatin A (20µM final concentration, Sigma). Chymotrypsin (Sigma) and trypsin (Sigma) were added at 14µg/ml final concentration. Following the proteolysis assay, samples were mixed with Laemmli sample buffer, heated in a thermal cycler to 95°C for 7 min., and subjected to SDS-PAGE for Western blotting.

### Western blot analysis of *in-vitro* proteolysis assays

Following the 24 hour incubation of reovirus and either T.E.E., or M.E.E (as mentioned above) at 37°C, 1x protein sample buffer (PSB; Laemmli buffer) was added to each sample, which was then heated to 95°C for 7-10 minutes and run on a 12% SDS-PAGE gel. A Bio-Rad Trans-blot Turbo system was used to transfer the SDS-PAGE gel to a nitrocellulose membrane (Amersham Hybond ECL, GE Healthcare Life Sciences). Following the transfer, the blots were then incubated overnight in 3% BSA/TBS-T (3% bovine serum albumin in tris-buffered saline with 0.1% Tween 20) blocking buffer. This incubation was followed by a 1 hour treatment with primary antibodies against reovirus capsid proteins (anti-reo; 1:1000 dilution in 3% BSA/TBS-T), reovirus core proteins (anti-core; 1:200 dilution) or the reovirus cell attachment protein σ1 (anti-σ1N(tail), anti-σ1C(head); 1:500 dilution). Primary antibodies (polyclonal mouse IgG) were kindly provided by Drs. Patrick Lee and Roy Duncan (Dalhousie University) and Dr. Terence Dermody (University of Pittsburgh). Goat anti-rabbit secondary antibodies conjugated with horseradish peroxidase (HRP; 1:10000 dilution) or Alexa Fluor-647 were used (1:5000 dilution) (Jackson ImmunoResearch). All antibodies were diluted in 3% BSA/TBS-T. Following application of each antibody, the blots were subjected to washing steps (totaling 15 minutes) prior to treatment with the next antibody or detection reagent. For all HRP antibodies, Pierce ECL (enhanced chemiluminescent) peroxidase substrate was used as a detection reagent. The blots were imaged and quantified using an ImageQuant LAS4000 biomolecular imager.

### Binding assay

HeLa, MDA-MB-231, L929, and MCF-7 cells were seeded in 100mm plates. Cells were grown until confluent, washed with PBS, detached from the plates with cell stripper, resuspended in PBS, then 5×10^5^ cells of each cell line were transferred to 1.5mL Eppendorf tubes. All steps of the binding assay were carried out at 4°C to allow binding but prevent internalization. Cells were incubated for 30 minutes, followed by infection for 1 hour on a rotator with the indicated reovirus variants Cells were centrifuged at 550 x g for 5 minutes at 4°Cand pellets were washed twice with 500µl 10% FBS in PBS solution. Final pellets were resuspended in 300µl of anti-reovirus primary antibody (probes for μ1C, δ and σ3, diluted 1:1000 in 10% FBS in PBS solution), and rotated for 45 minutes. Cells were then centrifuged as above, washed twice with 700ul 10% FBS in PBS, resuspended with 300µl of anti-rabbit AF-647 secondary antibody (Jackson Immunoresearch, 111-495-144, diluted at 1:500 in 10% FBS in PBS), and rotated for 45 min at 4 °C. Cells were then washed as above following primary antibody treatment. Final pellets were resuspended in 350µl of 4% paraformaldehyde (Sigma) and incubated for 45min. Cells were then centrifuged at 1000 x g for 7 minutes, washed with cold 1x PBS and resuspended in 300µl PBS. Cells were then analyzed on the BD LSRFortessa and 10,000 cells gated using BD FACS Aria III for viable and singlet cells based on FSC and SSC parameters. Mean fluorescence intensity was compared for samples exposed to virus relative to mock (PBS)-exposed cells, and presented without further gating strategies. To compare relative binding efficiencies between samples, background MFI (mock) was always subtracted from remaining MFI values. Then, a standard curve of particle dilution versus MFI was generated for positive control (wild-type T3D and corresponding cell line). MFI was then used to determine relative bound virus particles for remaining samples, keeping in mind that cells were exposed to equivalent virus particles of all treatments and variants in a given experiment. FCS Express 5 and FLOWJO software programs were both used to present data. For binding assays with MCF7, MTHJ, TUBO and T47D cells, a similar protocol was used with the following changes: 2×10^5^ cells were used, washing was completed in 1ml 2% FBS in PBS with 1uM EDTA, anti-sigma primary antibody (hybridoma bank 4F2) and secondary goad anti-mouse alexa 647 were used for detection of cell-bound virions.

### Immunocytochemistry

Cells were seeded in 24 well plates and infected with reovirus (MOI as indicated) at confluence, followed by incubation for 1 hour at 37 °C. Virus was removed by aspiration, and replaced with fully supplemented MEM. Cells were incubated for 12 hours at 37 °C, then fixed with methanol for 5 minutes and washed with PBS. Cells were then blocked with 1mL of blocking solution (PBS, 0.1% triton X100, 3% BSA). After 30 minutes blocking, solution was aspirated and 400µl of rabbit anti-reovirus primary antibody (diluted 1:10000 in block solution) was added to each well, and incubated for 1hr at room temperature. After removing the antibody, cells were washed 3x with wash solution and incubated with 400µl of alkaline phosphatase conjugated secondary antibody (Jackson Immunoresearch, “111-055-144”) (diluted 1:5000 in blocking solution) for 1hr. After discarding the secondary antibody, cells were rinsed with alkaline phosphatase buffer (10mM Tris, pH 9.5), and 400ul BCIP/NBT (5-bromo-4-chloro-3’-indolyphosphate/nitroblue tetrazolium, BCIP-B8503-500-Sigma, NBT-N6639-1G-Sigma) was added to each well. The cells were incubated in the dark at room temperature for 10-30 minutes. The reaction was stopped with a stop solution (10mM EDTA in PBS) and imaged on the EVOS Cell Imaging Systems (Invitrogen, Thermo Fisher Scientific).

### SNA staining and neuraminidase treatments

MCF7, MTHJ, TUBO, and T47D cells were grown until confluent, washed with PBS, detached with cell stripper, resuspended in PBS, then 6×10^5^ cells of each cell line were added. Cells were centrifuged at 350 x g for 5 min and pellets were washed twice with 1mL flow buffer (0.1% BSA in HBSS 1X +magnesium chloride +calcium chloride), after which they were incubated with 120mU neuraminidase (Sigma, from Clostridium perfringens) in PBS at 37°C for 30min. Cells were then centrifuged twice with 1mL flow buffer at 350 x g for 5min, and resuspended in 50ul of fluorescein labeled Sambucus nigra lectin (Vector Laboratories, SNA diluted at 1:1000 in washing solution) in the dark for 20min. All samples were fixed with 4% paraformaldehyde for 30min, centrifuged at 700 x g for 3min, and resuspended in flow buffer. Binding of SNA to sialic acid on cells was analyzed by flow cytometry.

### *In vivo* tumor model

Animal experiments were conducted with the approval of the University of Alberta Health Sciences Animal Care and Use Committee in accordance with guidelines from the Canadian Council for Animal Care. NOD-*scid* IL2Rgamma^null^ (NSG) mice were bred in-house by Dr. Lynne Postovit. Mice were housed in a biosafety level 2 containment suite at the University of Alberta Health Sciences Laboratory Animal Services Facility. Six-week-old NSG mice were implanted with 17β-estradiol pellets (0.72 mg/pellet 60-day release, Innovative Research of America, Cat. No. SE-121) at the back of the neck One week later, mice were sorted to give an even distribution of tumor sizes between groups, then MCF-7 cells (2 × 10^6^ cells in 50 μL 50% Matrigel) were injected into the mammary fat pad. When tumors were palpable, mice were injected 3 times with PBS or 10^6^ PFU of T3D, T3D(T249I), T3D(S18I) and T3D(S18I/T249I) viruses in 50µl. After virus injections, mice were monitored twice daily and weighed every other day. Tumor growth was measured by caliper twice weekly. Mice were euthanized by CO_2_ inhalation when virus treated animals presented initial symptoms of black foot/tail syndrome. Tumor and heart excised and frozen at -80°C. To determine titers, samples were thawed, weighed and resuspended in 2mL of HBSS buffer (ThermoFisher Scientific, Cat. No. 14025092). Next, samples were homogenized with a VDI 25 S41 homogenizer (VWR, Cat. No. 03231116), using 3 pulses of 10 seconds at highest speed (24.000 rpm). Samples were titered in L929 cells by standard plaque assay (70).

### Statistical analysis

Statistical methods are described in figure legends for respective experiments. Statistical analysis was performed using GraphPad Prism Version 7.05.

## ACKNOWLEDGEMENTS

We thank Dr. Matthew Macauley (University of Alberta) for providing expertise on sialic acids and the protocol for SNA staining. We thank Dr. Richard Schulz (University of Alberta) for providing expertise on metalloproteases. This work was funded by a Cancer Research Society project grant to MS, a grant to MS and MH from the Canadian Cancer Society, a project grant from the Li Ka Shing Institute of Virology to MS and MH, a salary award to MS from the Canada Research Chairs (CRC) and infrastructure support from Canada Foundation for Innovation (CFI), along with salary support for HE from a Canadian Institutes of Health Research (CIHR) project grant to MS. JF and SH received stipend funding from the University of Alberta Undergraduate Research Initiative (URI), and JF and IB also received stipend funding from the Alberta Innovates Health Solutions (AIHS). FC received stipend funding from the University of Alberta Faculty of Medicine & Dentistry, the John & Rose McAllister Graduate Scholarship, the Faculty of Graduate Studies & Research at the University of Alberta, and the John Thibault Memorial Fund. Flow cytometry was performed at the University of Alberta, Faculty of Medicine and Dentistry Flow Cytometry Facility, which receives financial support from the Faculty of Medicine and Dentistry and Canadian Foundation for Innovation (CFI) awards to contributing investigators.

